# A new insight into the role of CART peptide in serotonergic function and anxiety

**DOI:** 10.1101/2024.01.22.576719

**Authors:** Nagalakshmi Balasubramanian, Ruixiang Wang, Shafa Ismail, Benjamin Hartman, Zeid Aboushaar, Catherine A. Marcinkiewcz

**Affiliations:** Department of Neuroscience and Pharmacology, University of Iowa, Iowa City, IA; Iowa Neuroscience Institute, University of Iowa, Iowa City, IA

**Author notes:** Indicates co-corresponding author Please send all correspondence to: Catherine A. Marcinkiewcz, Department of Neuroscience and Pharmacology University of Iowa, Iowa City, IA-52242, USA.

**Keywords:** CART, 5HT, Anxiety, Dorsal raphe nucleus, Edinger Westphal nucleus

## Abstract

Cocaine and amphetamine-regulated transcript (CART) peptide has been established as a contributor to anxiogenic behavior. Genetic mutations in the CART gene are associated with anxiety and depression, and increased CART expression has been reported in suicide victims. Extensive research has focused on the role of CART peptide in mesolimbic neurocircuitry, but its involvement in the dorsal raphe nucleus (DRN) and serotonin (5HT) system remains unexplored. Here we demonstrate that CART processes are proximal to 5HT^DRN^ neurons and that microinjection of CART _(55-102)_ peptide into the DRN has an anxiogenic effect in mice. Furthermore, central CART administration reduced cfos activation in 5HT neurons of the ventral DRN, which is a putative reward/anti-stress circuit. The inhibitory effect of CART on 5HT^DRN^ neuronal function and local 5HT release is further demonstrated with *in vivo* fiber photometry coupled with calcium and 5HT biosensors and by mass spectrometry. Moreover, using Cre-dependent retrograde tracing, we observed DRN-projecting CART neurons in the Edinger Westphal nucleus (EW), nucleus accumbens (NAc), and various hypothalamic nuclei including the ventromedial hypothalamus (VMH). Interestingly, based on *ex vivo* electrophysiological recordings, acute stress increased excitability in DRN-projecting CART neurons located in the EW, but not in the VMH or NAc. This suggests that the stress may promote anxiety-like behavior by activating the EW^CART^→5HT^DRN^ circuit that ultimately inhibits 5HT transmission. In sum, understanding the intricate dynamics of the CARTergic and 5HTergic systems proves crucial in addressing 5HT-related dysfunctions, providing invaluable insights into both health and disease.

## Introduction

Approximately 280 million individuals globally are affected with anxiety and depression, a number that has surged by 25% since the COVID-19 pandemic [1]. Neuropeptides are neuromodulators that are frequently co-released with neurotransmitters and have been implicated in anxiety and depression [2]. Emerging evidence indicates cocaine and amphetamine-regulated transcript (CART) as a key player in regulating emotion, with a missense mutation in the CART gene being associated with heightened anxiety/depression in an adolescent population [3]. Moreover, an increased CART expression in the Edinger Westphal (EW) nucleus has been observed among suicide victims [4].

CART neurons implicated in stress response are predominantly localized in the arcuate nucleus (ARC), nucleus accumbens (NAc), EW, ventral tegmental area, locus coeruleus, and sparsely within the amygdala and hippocampus (HP) [5,6]. Intracerebroventricular (ICV) administration of CART _(55-102)_ into either the lateral ventricle (LV), NAc, or amygdala produces anxiogenic effects [7–9], whereas heightened CART levels in the HP, ARC, and paraventricular hypothalamic nucleus (PVN) were observed in stressed rats [10–12]. Furthermore, central administration of CART increased c-fos expression in corticotropin-releasing factor (CRF) neurons of the PVN, thereby modulating CRF secretion and underscoring CART’s crucial role in mediating the stress response [13].

While the regulatory influence of CART in anxiety, depression, and stress within the mesolimbic circuitry has been characterized, its physiological role in the brainstem nuclei, particularly the dorsal raphe nucleus (DRN) which is a central hub for serotonin (5HT) neurons, remains unexplored. Given the substantial contribution of 5HT in mood regulation [14–16], we aim to investigate the interplay between the CARTergic and 5HTergic systems, building upon our preliminary observations of CART projections to 5HT^DRN^ neurons. To ascertain the functional role of CART in the DRN, we investigated the impact of CART microinfusion on anxiety-like phenotypes in C57BL/6J mice. Employing state-of-the-art techniques, we explored the physiological and functional aspects of CART neurons and their role in modulating 5HT^DRN^ neuronal activity and anxiety.

## Methods

### Animals

Adult male C57BL/6J (000664), Fos2A-iCreER (030323), Sert-Cre (014554), Ai14 (007908), and Cart-IRES2-Cre-D (028533) mice from Jackson Labs (Bar Harbor, ME, USA) were used in these experiments. All mice were kept in a temperature- and humidity-controlled vivarium with *ad libitum* access to food and water under a 12h/12h light/dark cycle in accordance with AAALAC guidelines. All experimental procedures were approved by the IACUC at the University of Iowa.

### Immunofluorescence

Immunofluorescence was performed to localize CART and TPH2 neurons in the DRN of C57BL/6J as described previously [17,18]. 4-5 DRN sections were used across the rostral-caudal axis. Sections were permeabilized in 0.5% Triton X-100/PBS for 30 min, followed by blocking in 10% NDS-0.1% Triton X-100-PBS solution, and then incubated with primary and secondary antibodies (Supplementary Table 1). Images were acquired on an Olympus FV-3000 confocal microscope.

### Stereotaxic surgeries

Stereotaxic surgeries were performed for single/bilateral/fiber-cannula implantations (Supplementary Table 2) and intracranial AAV infusions. Surgeries were performed using an Angle Two stereotaxic frame (Leica Biosystems, Wetzlar, Germany). 250 nl of AAVrg-hsyn-DIO-EGFP (Addgene # 50457; MA, USA) was infused into the DRN (ML = ±0.0, AP = −4.65, DV = −3.30, θ = 23.58°) with a Quintessential Stereotaxic Injector (Stoelting, IL, USA) at a rate of 50 nl/min. For fiber photometry experiments, a fiber-cannula (OmFC) was implanted in the DRN and AAV was infused at the rate of 100 nl/min through OmFC guide cannula into the DRN using a fluid injector. Diluted 500 nl of AAV1-CAG-Flex-GCaMP6s-WPRE-SV40 (Addgene # 100842) and AAV9-syn-5HT3.5 (WZ Biosciences, MD, USA) was infused into the DRN of Sert-cre and C57BL/6J mice respectively.

A single cannula (P1 technologies, TX, USA) was implanted into the DRN of C57BL/6J for CART microinjection. In addition, bilateral cannulae were implanted in the LV (ML = ±1.0, AP = −0.30, DV = −2.20) of Fos2A-iCreER mice for experiments involving targeted recombination in activated populations (fos-TRAP).

### Drugs

Lyophilized CART peptide _(55-102)_ (003-62, Phoenix Pharmaceuticals, CA, USA) was reconstituted in 0.2 M PBS to obtain a stock concentration of 1 ug/ul. The final working CART peptide solution was freshly made in aCSF. For TRAP experiments, 4-hydroxytamoxifen (4-OHT; H6278; Millipore Sigma, MA, USA) was made as described previously [19]. Briefly, 20 mg/ml 4-OHT stock solution was prepared in ethanol by incubating at 40°C for 2-3 min. A working concentration of 5 mg/ml was made consisting of a 1:4 mixture of castor and sunflower seed oil. Ethanol from the working solution was evaporated by heating at 56 °C. Mice were injected at a volume of 10 ml/kg for a final dose of 50 mg/kg (i.p.).

### Behavioral tests

All behavioral tests were performed in C57BL/6J mice and recorded after 15 min of CART/aCSF infusion as reported previously [20,21]. Before the test day, the mice were habituated to infusion to reduce handling stress. CART peptide (5 ng, 50 ng, and 100 ng) or aCSF was infused through the guide cannula into the DRN.

#### Elevated plus maze test

Anxiety in the microinjected CART mice was assessed in the elevated plus maze (EPM) as reported previously [22]. After aCSF/CART infusion, mice were placed in the center of an elevated plus-shaped maze with white open and dark/black closed arms and allowed to explore for 5 min. Sessions were recorded with a Basler GenICam and Media Recorder software (Noldus, VA, USA). Time spent in open and closed arms and the number of entries to each arm were analyzed using EthoVision XT15.

#### Light-Dark box exploration test

The light-dark box exploration (LDB) test was conducted in the aCSF/CART microinjected mice as reported previously [23]. The apparatus consists of a light (390 lux) and a dark (5 lux) compartment. After infusion, the mice were placed in the center of the light compartment and the behavior was recorded for 10 min with a camera and Media Recorder software (Noldus, VA, USA). The time spent in each compartment was analyzed with EthoVision XT15.

#### Social interaction test

Social deficits were measured in the aCSF/CART microinjected mice in the social interaction test (SIT) as described previously [17]. Briefly, the test was conducted in a three-chambered Plexiglass with a small opening between the chambers. The test mouse was habituated to the entire arena for 10 min. At the end of the habituation phase, the test mouse was confined to the center chamber while a conspecific stranger mouse was placed inside a metal holding cage in one of the outermost chambers and an empty holding cage was placed in the opposite chamber. The social interaction test began by removing the doors between the chambers so that the test mouse could move freely throughout the entire arena for another 10 min. The behavior was recorded, and the time spent by the test mouse interacting with the conspecific stranger mouse and the empty cage was scored by a trained observer who was blinded to the experimental treatment.

### Targeted Recombination in Transiently Active Populations

To label neurons selectively activated by CART, we employed the TRAP method as described previously [19]. Before the test day, the mice were habituated to i.p injections and infusions to minimize handling stress. On the test day, either aCSF or CART (0.5 ug/mouse) was bilaterally infused in the LV of Fos-icreER x Ai14 mice. These mice express tdTomato in a Cre-dependent manner after induction of cfos. After 30 min of infusion, 4-OHT (50 mg/kg, i.p) was administered, and the mice were left undisturbed for several hours. After two weeks, the brains were extracted, and IF was performed on DRN slices with TPH2 antibodies. The images were captured on a confocal microscope and analyzed for colocalization of tdTomato and TPH2 signals using QuPath software (Version 0.4.2).

### Liquid Chromatography Mass Spectrometry (LCMS)

Neurotransmitter levels in the DRN after CART/aCSF infusion were measured using LC-MS. After 15 min of aCSF/CART microinjection into the DRN, the brains were extracted and snap-frozen immediately. DRN tissue was micro-punched, and LC-MS was performed at the University of Iowa Metabolomics Core. The samples were processed as described previously [24], and the data were acquired on a Thermo Q Exactive hybrid quadrupole Orbitrap mass spectrometer (Thermofisher Scientific, MA, USA). Millipore SeQuant ZIC-pHILIC (2.1 × 150 mm, 5 µm particle size) with a ZIC-pHILIC guard column (20 x 2.1 mm; Millipore-Sigma, MA, USA). MS data acquisition was performed in the range of m/z 70–1,000 with the resolution set at 70,000, the AGC target at 1×106, and the maximum injection time at 200 ms [24]. LC-MS data were processed by Thermo Scientific TraceFinder 4.1 software, and metabolites were identified based on an in-house library. NOREVA software was used for signal drift correction [25]. Data were normalized to the sum of all the measured metabolite ions in that sample.

### Fiber Photometry

Home cage fiber photometry experiments were performed to investigate the effects of CART on 5HT neuronal activity and transmitter release. Mice were habituated to patch cord tethering and infusion for 3 days. Data were recorded using a fiber photometry system (Neurophotometrics, CA, USA).

#### GCaMP6s biosensor in 5HT^DRN^ neurons

The effect of CART on 5HT neuronal activity was measured in Sert-cre mice injected with AAV1-CAG-Flex-GCaMP6s-WPRE-SV40. Signals were recorded from alternating pulses of 470nm (calcium-dependent) and 415nm (calcium-independent, isosbestic) with an excitation power set at 60 µW. Fiber photometry data were acquired with Bonsai software (Open Ephys, GA, USA) and processed via MATLAB R2021b software (MathWorks, MA, USA) as described previously by our group [26]. For all recordings, 470 and 415 signals were deinterleaved and processed to create a linear regression model using the MATLAB *fitlm* function to fit the raw data and correct motion artifacts. Next, each fluorescent intensity value was normalized to the corresponding value predicted by the linear regression model. Finally, we computed peri-event fluorescent changes (ΔF/F) between the last 5 min pre-infusion event (baseline) and 12 min post-infusion event and compared the area under the curve (AUC) in the peri-event plot for the CART versus aCSF groups.

#### 5HT3.5 biosensor in DRN

The effect of CART on 5HT release was measured in C57BL/6J mice injected with AAV9-hsyn-5HT3.5 (GRAB_5-HT_). For 5HT3.5 recordings, 415 isosbestic signals were not used in the analysis as it is not an appropriate wavelength for motion control for this sensor [27]. Peri-event fluorescent plots were generated by normalizing traces to the first 20s of each trace.

### Retrograde Tracing

DRN–projecting CART neurons were identified by a cre-dependent retrograde tracing approach as reported previously [28]. AAVrg-hsyn-DIO-EGFP was injected in the DRN of CART-cre x Ai14 reporter mice. These mice express tdTomato in a cre-dependent manner in CART neurons across the brain. After three weeks of viral expression, brains were extracted for analysis.

CART-expressing brain regions, especially those involved in stress and anxiety-related responses, were included in this study [5]. NAc, EW, medial amygdala (MeA), hypothalamic nuclei– ventromedial hypothalamus (VMH), tubular lateral hypothalamus (TuLH), ARC, PVN, and HP are the key targets we chose for retrograde tracing. The images were captured on a confocal microscope and analyzed for colocalization of tdTomato (CART neurons) and EGFP (DRN-projecting CART neurons) in QuPath software (Version 0.4.2).

#### *Ex vivo* electrophysiology

Three weeks before the experiments, CART-Cre x Ai14 mice were injected with AAVrg-hsyn-DIO-EGFP in the DRN for retrograde tracing. Restraint stress (30 min) was used to mimic the anxiety-like response evoked by the behavior tests, and undisturbed mice served as controls. The method of stress induction prior to *ex vivo* electrophysiological recordings has been described in detail previously [29]. Acute 300-µm brain slices containing the EW, VMH, and NAc were prepared. Fluorescent filters were utilized to screen cells expressing both tdTomato and EGFP, i.e., DRN-projecting CART neurons. Patch electrodes were filled with a solution containing (in mM): 135 K-gluconate, 5 NaCl, 2 MgCl_2_, 10 HEPES, 0.6 EGTA, 4 Na_2_-ATP, and 0.4 Na_2_-GTP. The current clamp mode was employed to assess intrinsic excitability, and cells were kept at −70 mV to offset the variation in resting membrane potential. Rheobase was measured by the minimal current needed to evoke action potentials. Input resistance was examined by the decrease in membrane potential after injection of a −100 pA hyperpolarizing current. The number of spikes (i.e., evoked action potentials) were recorded upon injection of depolarizing currents for 250 ms at 10 pA incremental steps (0 – 200 pA).

### Data and statistical analysis

Statistical analysis was performed using GraphPad Prism 10 (CA, USA). Student *t-*tests were used for comparisons between two groups, and a one-way or two-way ANOVA was applied for comparisons between more than two groups with one or more independent variables respectively. A *p*-value less than 0.05 was considered significant for all the analyses. Data are expressed as mean ± SEM.

## Results

### Microinfusion of CART peptide in the DRN induces anxiogenic behavior

Immunofluorescence experiments revealed that CART axonal terminals are proximal to 5HT neurons, suggesting that CART peptide may be released in the DRN and alter behavioral processes by modulating the activity of 5HT neurons (Fig. 1A, B). We then aimed to determine if CART could elicit anxiogenic effects upon injection into the DRN of male C57BL/6J mice (Fig. 1C). Various CART doses (5 ng, 50 ng, and 100 ng) were microinjected into the DRN, and anxiety-like behaviors were assessed using the EPM, LDB, and SIT. Mice receiving a high CART dose (100 ng) exhibited heightened anxiety in the EPM, spending more time in the closed arm (one-way ANOVA, *F*_3,30_ = 4.7984, *p* < 0.001, followed by *post hoc* Dunnett’s tests showing aCSF vs 100 ng: *p* < 0.01) versus the open arm (one-way ANOVA, *F*_3,30_ = 4.154, *p* < 0.05, followed by *post hoc* Dunnett’s tests showing aCSF vs 100 ng: *p* < 0.001) (Fig. 1D-E). Also, a non-significant trend towards increased time in the closed arm over the open arm in the EPM was observed in the two lower dose (5 ng and 50 ng) groups. Additionally, the CART-100 ng group showed a significantly reduced number of entries into the open arm (one-way ANOVA, *F*_3,30_ = 3.250, *p* < 0.05, followed by *post hoc* Dunnett’s tests showing aCSF vs 100 ng: *p* < 0.05) (Fig. 1F-G). The heat-map visualization of the EPM test illustrates a CART dose-dependent increase in anxiety in comparison to aCSF controls (Fig. 1H-K). The LDB test exhibited a similar dose-dependent anxiogenic effect with the highest dose of 100 ng showing the strongest anxiety effect compared to lower doses and the vehicle controls. Mice receiving 100 ng of CART spent more time in the dark box (one-way ANOVA, *F*_3,28_ = 3.538, *p* < 0.05, followed by *post hoc* Dunnett’s tests showing aCSF vs 100 ng: *p* < 0.05) versus the light box (one-way ANOVA, *F*_3,28_ = 3.808, *p* < 0.05, followed by *post hoc* Dunnett’s tests showing aCSF vs 100 ng: *p* < 0.05) (Fig. 1L-M). Next, we measured social deficits in the SIT and observed a decrease in the percentage of time spent in social interaction in the CART-100 ng group as compared to aCSF controls (one-way ANOVA, *F*_3,30_ = 3.129, *p* < 0.05, followed by *post hoc* Dunnett’s tests showing aCSF vs 100 ng: *p* < 0.05). Interestingly, the time spent interacting with the empty cage (EC) increased, while interaction with a stranger cage (SC) decreased in the CART-100 ng group compared to the aCSF group (two-way ANOVA: interaction effect between aCSF/CART and EC/SC, *F*_1,28_ = 12.07, *p* < 0.01, with a *post hoc* uncorrected Fisher’s LSD test showing increased EC interaction, *p* < 0.05 and decreased SC interaction, *p* < 0.05 in mice infused with CART vs aCSF) (Fig. 1N-O). Collectively, these experiments indicate that intracranial infusion of CART into the DRN has an anxiogenic effect in mice.

**Figure 1:**
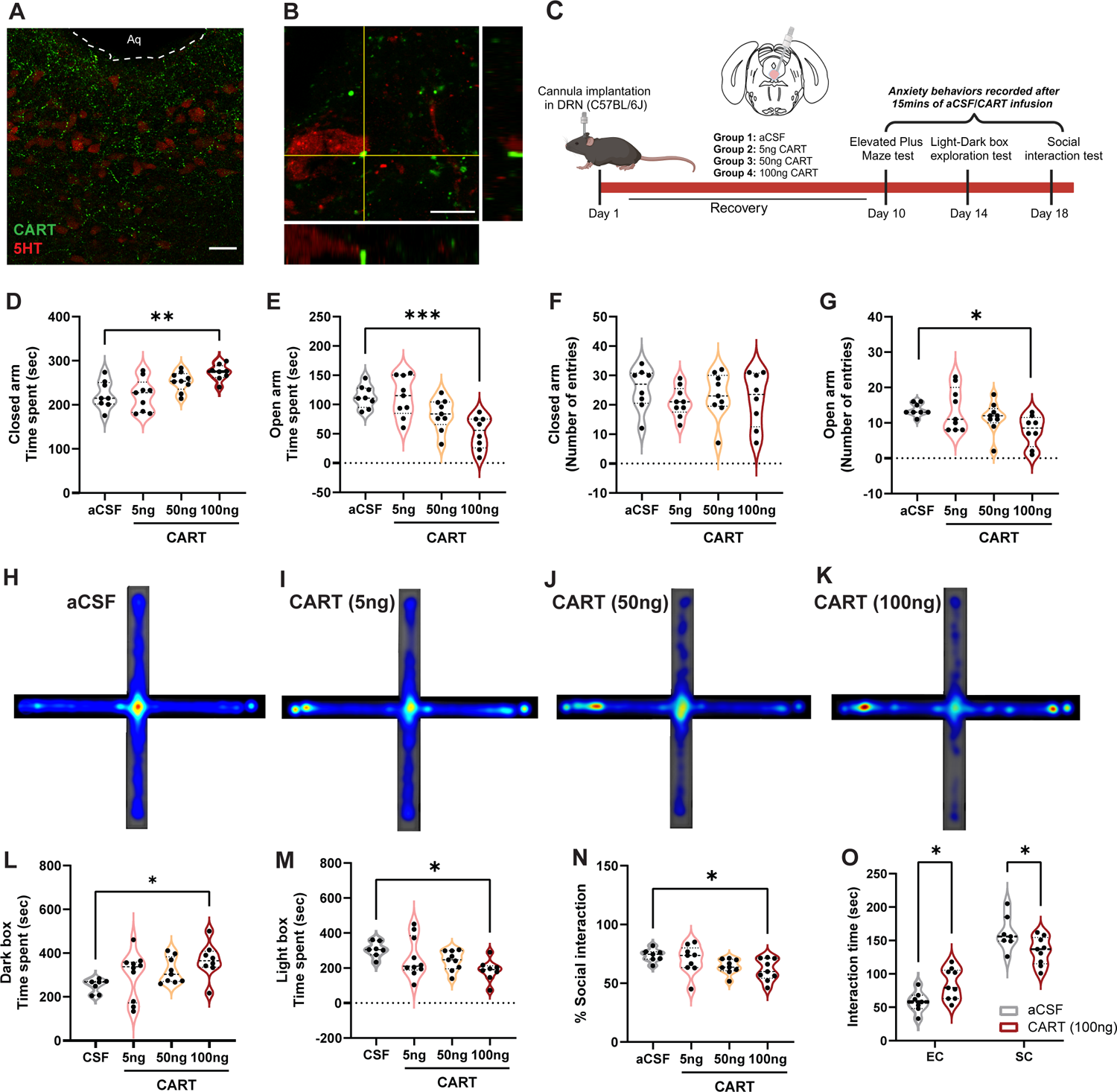
CART peptide infusion into the DRN increases anxiogenic behavior. (A) Representative confocal images of CART peptide and 5HT in DRN of C57BL/6J. Scale bar = 50 µm. (B) High-resolution orthogonal view showing the proximity of CART axon terminals to 5HT neurons. Scale bar = 10 µm. (C) Schematic representation of the experiments showing the surgery and behavior schedule. Elevated plus maze (EPM) test showing time spent in (D) closed arm; (E) open arm and (F-G) number of entries into each arm after CART infusion into DRN; (H-K) Heat map visualization of time spent by mice in the EPM arms that received either aCSF or CART (5 ng, 50 ng, 100 ng). Light Dark box exploration test showing time spent in (L) Dark box and (M) Light box in aCSF and CART microinjected mice. Social interaction test showing (N) % social interaction with strangers and (O) Interaction time with empty and stranger cages in aCSF and CART microinfused mice. Values (n = 7-10/group) are represented as means (±SEM) and the data was analyzed by One-way ANOVA (\**p* < 0.05, \*\**p* < 0.01, \*\*\**p* < 0.001 versus aCSF). *Part of the figure was made in BioRender*.

### ICV administration of CART peptide reduced fos activity in ventral 5HT^DRN^ neurons

Subsequently, we sought to investigate whether the anxiogenic effects mediated by CART were indicative of altered activity in DRN neurons. To address this, we conducted a fos-TRAP experiment utilizing Fos2A-iCre x Ai14 mice to assess CART-mediated 5HT neuronal activation (Fig. 2A). Microinjection of CART into the LV resulted in a non-significant decrease in cfos activity across the entire DRN as indicated by tdTomato-positive neurons (t_9_ = 1.398, ns; Fig. 2B-C). We then analyzed specific subregions and found that the rostral DRN region exhibited a significant reduction in tdTomato^+ve^ neurons (t_9_ = 2.704, *p* < 0.05), with no observable change in mid and caudal regions (Fig.2D). It was recently reported that 5-HT^DRN^ neurons differentially modulate stress-related behavior depending on their relative position across the dorsal-ventral axis, with neurons in the dorsal DRN activating subcortical stress pathways and neurons in the ventral DRN activating cortical anti-stress systems [30]. Therefore, we performed a separate analysis of the dorsal and ventral regions of the DRN and observed a significant decrease in tdTomato^+ve^ neurons in the ventral DRN upon ICV infusion of CART (t_9_ = 2.963, *p* < 0.05; Fig. 2E). Furthermore, a specific examination of cfos activation in 5HT^DRN^ neurons revealed a significant decrease in tdTomato^+ve^ 5HT neurons in the CART-100 ng compared to the aCSF control group (t_9_ = 2.943, *p* < 0.05; Fig. 2F-G), and the ventral 5HT neurons in particular displayed significantly lower tdTomato signal (t_9_ = 4.146, *p* < 0.01; Fig. 2H) upon CART infusion. Analysis of the proportion of tdTomato^+ve^ 5HT neurons within the 5HT neuron population indicated an overall decrease in cfos activity in 5HT neurons (t_9_ = 3.343, *p* < 0.05), with a significant reduction observed in mid (t_9_ = 2.869, *p* < 0.05), caudal (t_9_ = 3.477, *p* < 0.01) and ventral 5HT^DRN^ neurons (t_9_ = 3.183, *p* < 0.05; Fig. 2I-K). Together, these findings suggest that CART may promote anxiety-like behavior by inhibiting cortical-projecting ventral 5-HT^DRN^ neurons that activate anti-stress pathways in the brain.

**Figure 2:**
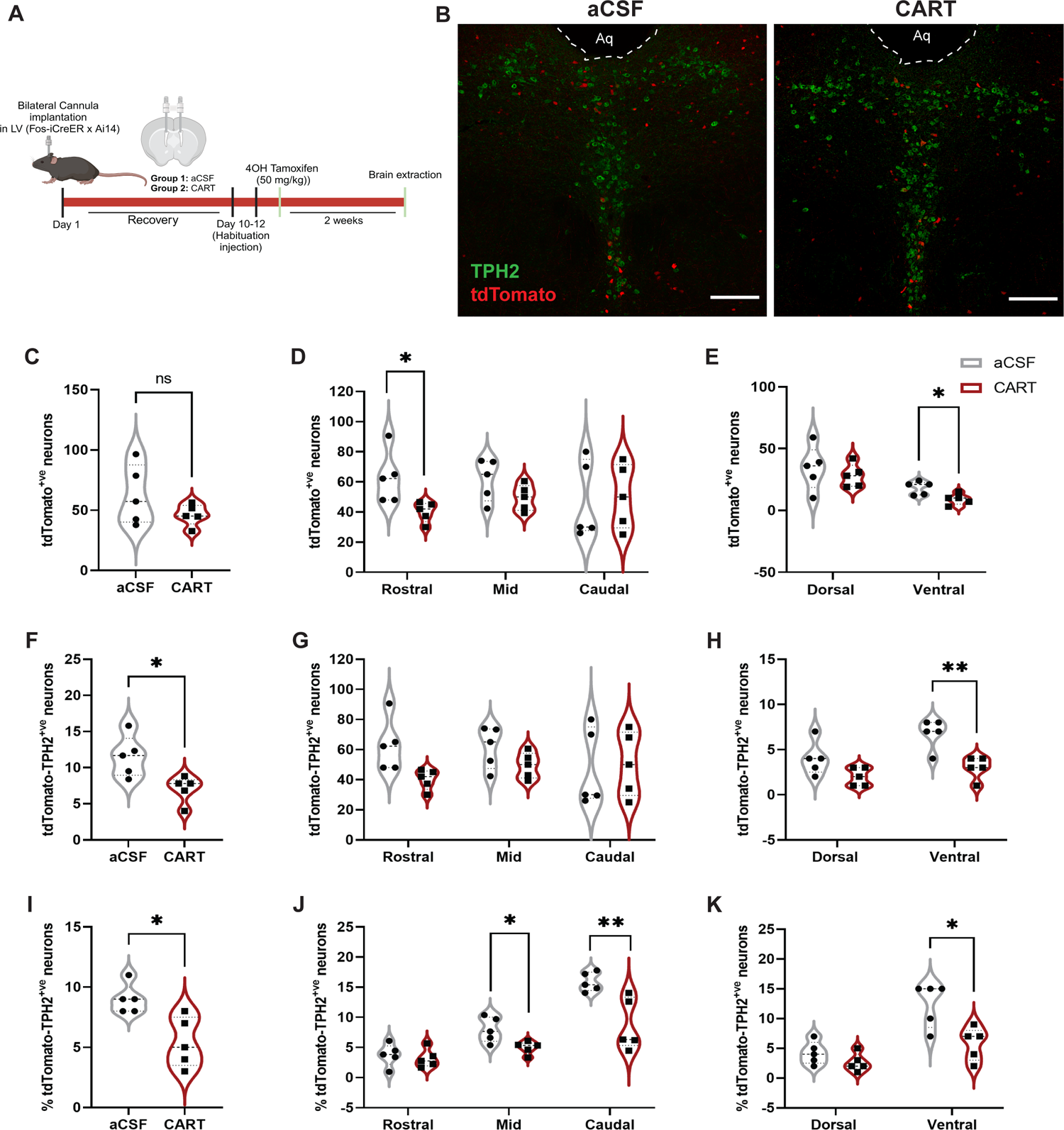
CART peptide decreases cfos induction in ventral DRN-TPH2 neurons. (A) Schematic representation of targeted recombination of active populations (fos-TRAP) experiments after CART infusion in the lateral ventricle (LV) of Fos-iCreER x Ai14 mice. (B) Confocal images showing tdTomato^+ve^ TPH2 neurons in aCSF and CART group. Scale bar = 200 µm. Violin plot showing the number of total tdTomato^+ve^ neurons in (C) DRN, (D) rostral, mid, and caudal regions, and (E) dorsal and ventral regions of DRN. Violin plot showing the number of total tdTomato^+ve^ TPH2 neurons in (F) DRN, (G) rostral, mid, and caudal DRN and (H) dorsal and ventral DRN. Violin plot showing the percentage of total TPH2 neurons positive for tdTomato signal in (I) DRN, (J) specific to rostral, mid, and caudal DRN and (K) dorsal and ventral DRN. Values (n = 5/group) are represented as means (±SEM) and the data was analyzed by single and unpaired t-test (\**p* < 0.05, \*\**p* < 0.01 versus aCSF). *Part of the figure was made in BioRender*.

### Microinfusion of CART peptide in the DRN reduces 5HT neuronal activity and transmitter release

#### LCMS

Next, we sought to elucidate the impact of CART on various neurotransmitters in the DRN. Micro-punched DRN tissues from mice that received CART/aCSF were processed for measuring neurotransmitter levels via LC-MS. Interestingly, LC-MS data revealed a significant decrease in serotonin levels in the CART-100ng vs aCSF group (one-way ANOVA, *F*_3,29_ = 2.497, *p* < 0.01, followed by *post hoc* Dunnett’s tests showing aCSF vs 100 ng: *p* < 0.05) (Fig. 3A). The chromatograph illustrating serotonin peak intensity indicated a decrease in 5HT levels upon CART infusion in the DRN (Fig. 3B). However, the 5HT metabolite, 5-Hydroxyindoleacetic acid (5HIAA), exhibited no significant change (Fig. 3C). Furthermore, GABA, glutamate, and HVA (Homovanillic acid, a dopamine metabolite) showed no apparent alterations following CART infusion (Fig. 3D-F).

**Figure 3:**
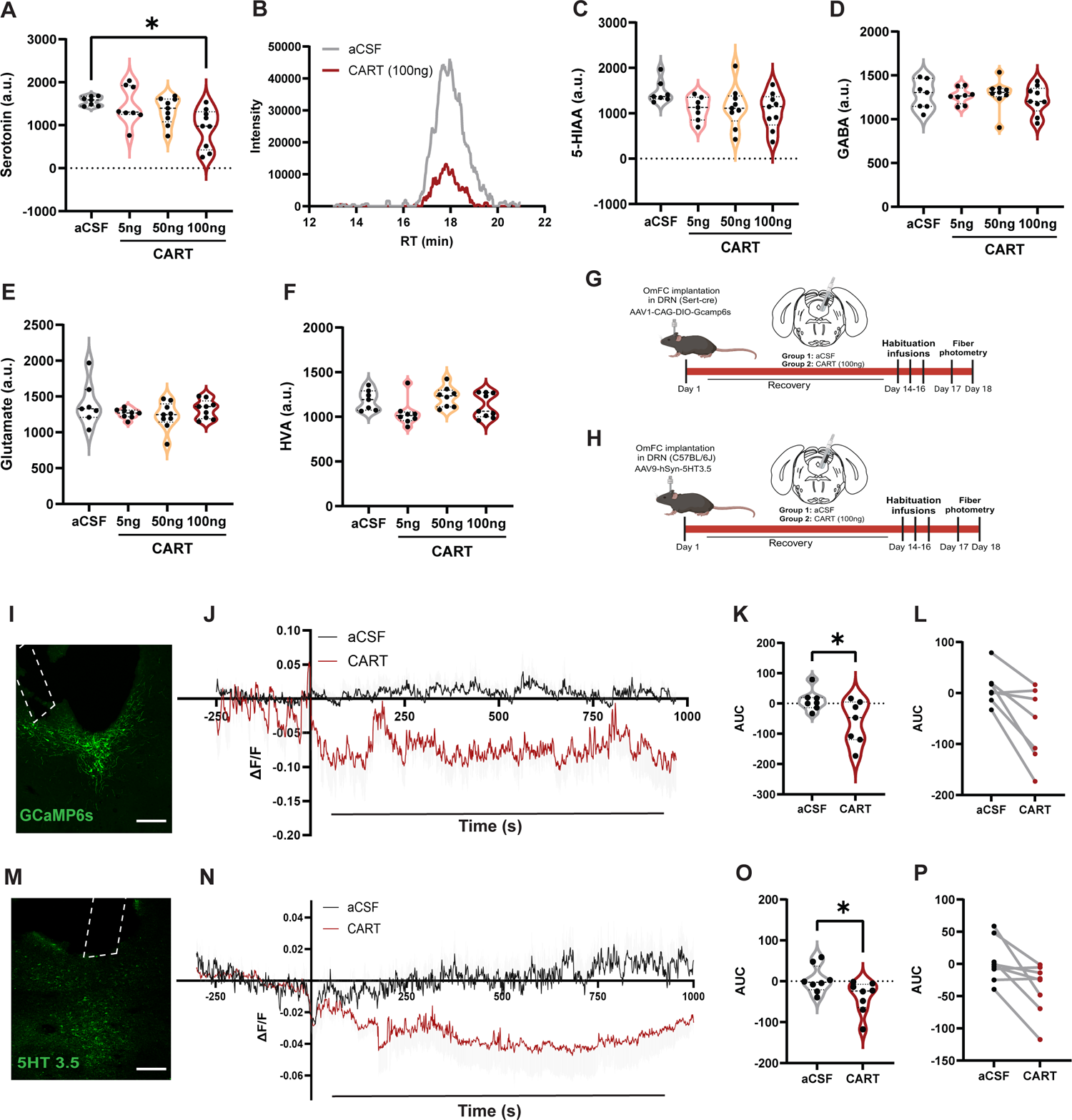
CART peptide inhibits 5HT neuronal activity and transmitter release. Violin plot showing (A) serotonin levels in DRN and (B) Chromatograph with decreased serotonin peak at 16-20 retention time (RT) in LC-MS after aCSF or CART (5 ng, 50 ng, 100 ng) infusion. Violin plot showing levels of (C) 5-hydroxy indole-acetic acid (5HIAA) (D) GABA (E) Glutamate (F) Homovanillic acid (HVA) in LC-MS after CART/aCSF infusion. Values (n = 8-10/group) are represented as means (±SEM) and the data was analyzed by one-way ANOVA and (\**p* < 0.05, versus aCSF). (G and H) Schematic representation of the timeline of the *in vivo* fiber photometry experiment using GCaMP6s and 5HT biosensors in Sert-Cre and C57BL/6J mice respectively. (I) Confocal image showing the expression of AAV1-CAG-Flex-GCaMP6s-WPRE-SV40 in DRN of Sert-Cre mice. Scale bar = 200 µm. (J) (ΔF/F) showing a drop in GCaMP signal as a measure of 5HT activity in the CART vs aCSF group. (K and L). Violin plot showing decreased area under curve (AUC) GCaMP values in the CART vs aCSF mice. (M) Confocal image showing the expression of AAV9-syn-5HT3.5 in DRN of C57BL/6J mice. Scale bar = 200 µm. (N) (ΔF/F) showing a dro*p* in 5HT3.5 signal as a measure of 5HT release in the CART vs aCSF group. (O and P). Violin plot showing decreased AUC values for 5HT3.5 in the CART vs aCSF mice. Values (n = 8/group) are represented as means (±SEM) and the data was analyzed by unpaired t-test (\**p* < 0.05, versus aCSF). *Part of the figure was made in BioRender*.

#### Fiber photometry

Next we monitored the impact of CART microinjection on 5HT^DRN^ neuronal activity in real time in awake behaving mice. For this experiment, Sert-cre mice were microinjected with a cre-dependent GCaMP6s calcium biosensor in the DRN and implanted with a specialized OmFC mount comprising an optical fiber for activity measurement and a cannula for aCSF/CART delivery (Fig. 3G and I). We observed a decrease in the GCaMP signal (ΔF/F) post-CART (100 ng) microinfusion into the DRN (Fig. 3J). This effect persisted for 15 min post-infusion, with significantly lower AUC values in the CART-100 ng group compared to the aCSF group (t_13_ = 2.400, *p* < 0.05) (Fig. 3K-L). Real-time recording of 5HT release in C57BL/6J mice using the 5HT3.5 biosensor corroborated the GCaMP results (Fig. 3H and M). The 5HT3.5 signal (ΔF/F) decreased after CART microinfusion, with concurrently reduced AUC values (t_15_ = 2.297, *p* < 0.05; Fig. 3O-P). Collectively, these findings suggest that CART diminishes 5HT^DRN^ neuronal activity and transmitter release, ultimately culminating in reduced 5-HT concentrations in the DRN. This complements our previous findings that CART reduces cfos activity in 5HT neurons in the ventral DRN, which were previously reported to promote stress-coping behaviors [30].

#### Retrograde tracing of CART projection neurons to the DRN

So far, the results suggest that CART microinjection can mimic endogenous CART release into DRN and may play a role in producing anxiogenic behavior via modulation of 5HT activity. We then sought to map the anatomical origin of these DRN-projecting CART neurons. To identify the specific population of DRN-projecting CART neurons, an AAVrg-hsyn-DIO-EGFP viral construct was injected into the DRN of CART-cre x Ai14 mice (Fig. 4A). Of the regions analyzed, the highest density of retrogradely-labeled CART neurons was observed in the EW (Fig. 4B and D), and the least was observed in the HP (Fig. 4B) and MEA (Fig. 4B and F). The EW showed the most intensely labeled CART neurons (Fig. 4B and E), which may reflect the fact that the EW is located proximal to the DRN. Various hypothalamic nuclei also contained labeled CART neurons that project to DRN. This includes the ARC (Fig. 4B and G), VMH (Fig. 4B and H), TuLH (Fig. 4B and I), and PVN (Fig.4B and J). The proportion of DRN-projecting CART neurons relative to the total CART population within these regions is depicted in Fig. 4C.

**Figure 4:**
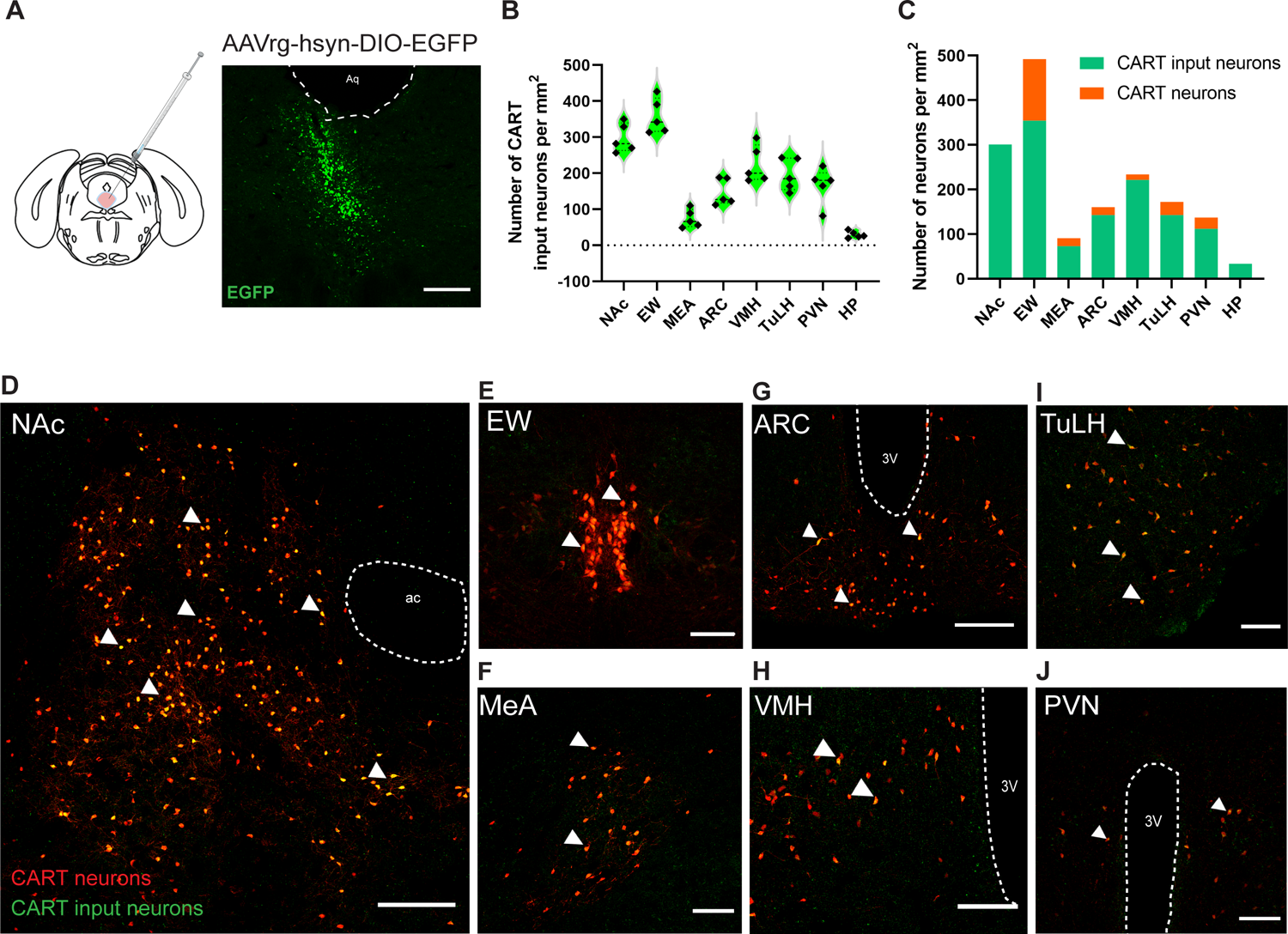
Retrograde tracing of DRN-projecting CART neurons. (A) Diagram showing the injection of a cre-dependent retrograde tracer (AAVrg-hsyn-DIO-EGFP) injected into DRN of CART-Cre x Ai14 mice and a confocal image showing the expression of EGFP after 3 weeks. Scale bar = 200 µm. (B) Number of CART positive retrograde labeled (input) neurons per mm^2^ in the nucleus accumbens (NAc), Edinger Westphal nucleus (EW), medial amygdala (MEA), arcuate nucleus (ARC), ventromedial hypothalamus (VMH), tubular hypothalamus (TuLH), paraventricular hypothalamus (PVN), and hippocampus (HP). (C) Proportion of CART-input neurons to total CART neurons in NAc, EW, MEA, ARC, VMH, TuLH, PVN and HP per mm^2^. Representative confocal image showing CART positive retrograde labeled neurons in (D) NAc (E) EW (F) MEA (G) ARC (H) VMH (I) TuLH and (J) PVN. Scale bar = 200 µm (NAc), and 100 µm (other regions). *Part of the figure was made in BioRender*.

#### Acute stress increased the excitability of DRN-projecting CART neurons located in the EW

Next, we used *ex vivo* slice electrophysiology to compare the intrinsic excitability of DRN-projecting CART neurons located in the EW, VMH, and NAc, three brain regions with the largest populations of such neurons. Interestingly, DRN-projecting CART neurons in the EW were the most excitable as assessed by rheobase (action potential threshold), input resistance, and evoked spike frequency, followed by those in the VMH and the NAc. Rheobase was highest in NAc neurons (183.80 ± 21.21 pA, *n* = 9), followed by VMH neurons (105.49 ± 24.89 pA, *n* = 8) and EW neurons (81.86 ± 10.27 pA, *n* = 8; one-way ANOVA, *F*_2,22_ = 7.39, *p* < 0.01). *Post hoc* Tukey tests confirmed that the rheobase of DRN-projecting CART neurons in the NAc was significantly higher (i.e. less excitable) compared to the other populations: EW vs. NAc: *p* < 0.01, VMH vs. NAc: *p* < 0.05, and EW vs. VMH: non-significant (Fig. 5A-B). In line with the results for rheobase, DRN-projecting CART neurons in the EW had the highest input resistance (302.5 ± 15.64 MΩ, *n* = 8), followed by VMH neurons (225.74 ± 28.28 MΩ, *n* = 9) and NAc neurons (113.40 ± 7.89 MΩ, *n* = 9; one-way ANOVA, *F*_2,23_ = 23.48, *p* < 0.0001, with *post hoc* Tukey tests showing EW vs. VMH: *p* < 0.05, EW vs. NAc: *p* < 0.0001, and VMH vs. NAc: *p* < 0.01 (Fig. 5C-D). Moreover, incrementally injecting depolarizing currents (at 20 steps, 10 pA/step) evoked more action potentials in EW neurons than in NAc neurons (two-way ANOVA: interaction effect between current and brain region, *F*_38,399_ = 2.66, *p* < 0.0001, with a *post hoc* Tukey test showing EW vs. NAc: *p* < 0.05; consistently, one-way ANOVA for AUC between the spike number curves and the x-axis: *F*_2,21_ = 3.89, *p* < 0.05, with a *post hoc* Tukey test showing EW vs. NAc: *p* < 0.05; (Fig.5E-G). In addition, hyperpolarization-activated cation current (Ih) appeared in every EW neuron we recorded, as shown by a prominent “sag” in voltage response to depolarizing current injection (Fig. 5C), which is consistent with a previous report [31].

**Figure 5:**
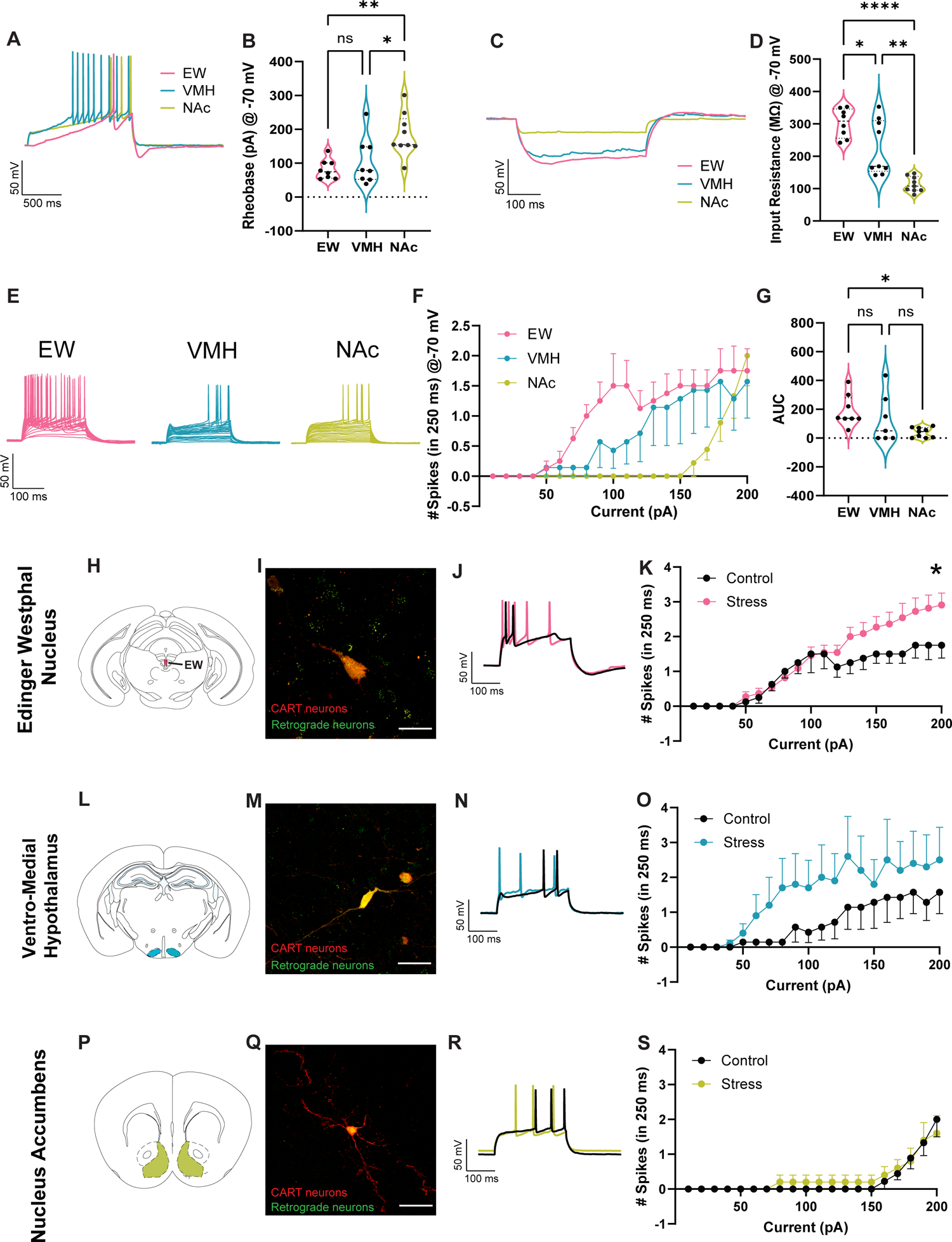
Acute stress increased the excitability of DRN-projecting CART neurons in the EW of CART-Cre x Ai14. (A) Representative traces for the rheobase experiments: incrementing the magnitude of injected currents until action potentials were evoked. (B) Histogram of rheobase in DRN-projecting CART neurons in the EW, VMH, and NAc when membrane potential was held at −70 mV. Those in the NAc had the greatest rheobase (i.e., minimal current to evoke action potentials). (C) Representative traces for the input resistance experiments: assessing changes in membrane potential upon the injection of −100 pA hyperpolarizing currents. (D) Violin plot of input resistance in DRN-projecting neurons in the EW, VMH, and NAc when membrane potential was held at −70 mV. Those in the EW had the highest input resistance, indicating the highest excitability. (E) Representative traces for assessing evoked spike (action potential) frequency upon the injection of incremental depolarizing currents for 250 ms at 20 steps (from 0 to 200 pA, with 10 pA/step). (F) Curves depicting numbers of spikes/250 ms evoked by incremental depolarizing currents in DRN-projecting CART neurons in the EW, VMH, and NAc when membrane potential was held at −70 mV. (G) Histogram of the area under the curve (AUC) shown in Panel F. AUC was greater for EW vs NAc, indicating higher excitability of EW neurons. (H), (L), & (P) Coronal sections that contain the brain regions targeted in electrophysiological recordings. (I), (M), & (Q) Confocal images of representative DRN-projecting CART neurons for recordings in the EW, VMH, and NAc, respectively. Scale bar = 20 µm (J), (N) & (R) Representative traces obtained at the 200 pA current step from the EW, VMH, and NAc, respectively. (K), (O), & (S) Curves depicting numbers of spikes/250 ms evoked by incremental depolarizing currents in mice with and without exposure to 30-min restraint stress from the EW, VMH, and NAc, respectively. Acute stress led to increased spike numbers in EW neurons. Data was analyzed by one-way and two-way ANOVA (\**p* < 0.05; \*\**p* < 0.01; \*\*\*\**p* < 0.001; ns: non-significant).

To assess whether DRN-projecting CART neurons in the EW, VMH, and NAc were associated with stress response and anxiety, we performed *ex vivo* electrophysiological recordings immediately after the mice were exposed to an acute (30 min) restraint stress. Our data showed that acute stress led to increased excitability of CART neurons in the EW, but not in the VMH or NAc [for evoked spike numbers, in the EW, two-way ANOVA: interaction effect between current and stress, *F*_19,323_ = 3.34, *p* < 0.0001, with a *post hoc* Šidák test showing that at the 200 pA current step, control vs. stress: *p* < 0.05, (Fig. 5H-K), and no significant interaction effects or main effects of stress were observed in the VMH (Fig. 5L-O) or NAc (Fig. 5P-S)]. Nevertheless, the high variability associated with DRN-projecting CART^VMH^ neurons in our recordings indicates that there could exist distinct neuronal subpopulations in this brain region, which warrants further investigation.

## Discussion

In the present study, we investigated for the first time the interactions between CARTergic and 5HTergic^DRN^ systems and their role in mediating anxiety states using state-of-the-art techniques including fiber photometry, LC-MS, fos-TRAP, and *ex vivo* electrophysiology. Our findings from high-resolution microscopy indicate prominent CART projections to 5HT^DRN^ neurons. Previous studies have shown that ICV administration and intracranial microinjection of CART in the amygdala or NAc induces anxiogenic behaviors in rats [7–9]. In the current study, we demonstrate that intracranial infusion of CART in the DRN produced an anxiogenic response (EPM and LDB) in mice. Moreover, CART microinfusion also induces social deficits in the SIT, and previously optogenetic stimulation of 5HT^DRN^ neurons has been demonstrated to rescue social deficits in Shank3 KO mice [32]. Our findings imply that CART may influence 5HT neurons to produce social deficits in the mice. This is further supported by the results of the fos-TRAP experiments wherein we demonstrate that CART inhibits 5HT^DRN^ neuronal activity in the ventral DRN, which is a subregion previously reported to promote stress-coping behaviors [30]. The reduced cfos activation in the DRN was accompanied by lower 5HT levels detected by LC-MS and reduced transmitter release measured by *in vivo* fiber photometry with a 5HT3.5 biosensor in freely moving mice. Inhibition of 5HT release by CART also paralleled the real-time reduction in calcium signaling in DRN-5-HT neurons in Sert-Cre mice.

Previous studies have localized DRN-projecting CART neurons in the ARC and PVN using red RetroBeads as a retrograde tracer [33,34]. Here we mapped DRN-projecting CART neurons in other nuclei–EW, ARC, VMH, LH, PVN, MEA, HP, neocortex, and NAc--using a Cre-dependent retrograde AAV (AAVrg) viral construct in CART-Cre x Ai14 mice, which allowed us to visualize both DRN-projecting and non-projecting CART neurons. CART is implicated in various functions including stress, anxiety, and depression[5]; and the DRN-projecting CART neurons are abundantly expressed in the nuclei that are known to regulate anxiety and stress responses [9,11,35,36]. *Ex vivo* electrophysiological recordings in DRN-projecting CART neurons in the EW, VMH, and NAc demonstrated different levels of intrinsic excitability at baseline with CART^EW◊DRN^ neurons as the most excitable. Moreover, acute restraint stress led to increased excitability only in CART^EW◊DRN^ neurons. In line with our findings, a recent report has shown that chemogenetic activation of CART^EW^ neurons increases anxiety while ablation or inhibition of CART neurons decreases anxiety [6]. However, many neuropeptides including nesfatin, pituitary adenylate cyclase-activating polypeptide, and cholecystokinin released by CART^EW^ neurons have been implicated in anxiety [37–39]. Although CART^EW^ neurons co-release other anxiety-regulating neuropeptides, the present study suggests that CART itself may be involved in exerting anxiogenic effects by inhibiting 5HT^DRN^ neurons. In support of our findings, higher EW-CART mRNA levels were observed in suicide victims [4].

We also observed high variability in basal excitability and stress responsiveness in DRN-projecting VMH neurons. Since hypothalamic CART neurons are heterogeneous in nature, they co-express other neurotransmitters that are crucial in appetitive or reward regulation [5,40]. Thus, all CART^VMH◊DRN^ neurons may not be involved in anxiety or stress responses, which needs to be examined in the future. Interestingly, DRN-projecting CART neurons in the NAc were the least excitable as compared to those located in the EW and VMH, and they showed no response to acute stress in *ex vivo* electrophysiology. The function of CART peptide in the NAc is not very clear but it has been suggested that it is a homeostatic regulator of dopamine-mediated activity, which may be excitatory or inhibitory [41]. However, in another study, stimulation of 5-HT_4_Rs in the NAc regulates anorexia by increasing CART mRNA levels [42]. There could be reciprocal projections between CART^NAc^ and 5HT^DRN^ neurons, but DRN-projecting CART^NAc^ neurons may be the primary mediators of stress or anxiety. Instead, they may play a significant role in regulating reward or feeding circuits [43].

Our study establishes for the first time the neuromodulatory role of CART in regulating 5HT neuronal function in the DRN. Our findings suggest that CART inhibits a specific sub-population of ventral DRN 5HT neurons associated with stress coping, consequently contributing to anxiety-like behavior. Endogenous CART originating in the EW has been previously implicated in anxiety and also projects to the DRN and is activated in response to acute stress. This strongly suggests that the CART^EW^-5HT^DRN^ circuit may be a key driver of anxiety-like behavior. Further investigation into the recently identified putative CART receptor, GPR-160, and its localization in this circuit will be essential to fully comprehend the inhibitory effects of CART on 5HT^DRN^ neurons.

## Acknowledgments

We thank Dr. Jeffrey Friedman (Rockefeller University, USA) for providing the Cart-IRES2-Cre-D mice for these studies. We acknowledge the University of Iowa Metabolomics Core Facility for performing LC-MS analysis. We thank Yu Xu for maintaining mouse colonies and Thomas James for technical assistance in fiber photometry.

## Author contributions

N.B and C.A.M. conceptualized the project and wrote the manuscript with contributions from all other co-authors. N.B. performed stereotaxic surgeries, behavioral experiments, retrograde tracing, fiber photometry experiments, and confocal imaging with the assistance of S.I. and B.H. R.W. performed whole-cell patch clamp electrophysiological recordings and analyzed the data with assistance from Z.A.

