## Supplementary tables for "A new insight into the role of CART peptide in serotonergic function and anxiety"

**Table 1: List of antibodies used in immunofluorescence.**

| **Stain** | **Antibody information** | **Concentration** |
| --- | --- | --- |
| Immunofluorescence CART-5HT | Goat anti-5HT  (Immunostar; catalog # 20079) | 1:2000 |
|  | Rabbit anti-CART  (Phoenix pharmaceuticals, catalog # H-003-062) | 1:2000 |
|  | 488 Donkey anti-Rabbit (Jackson ImmunoResearch; catalog # 711-545-152) | 1:500 |
|  | Cy3 Donkey anti-Goat  (Jackson ImmunoResearch; catalog # 705-165-147) | 1:500 |
| Immunofluorescence TPH2 (fos-TRAP) | Goat-TPH2 (Everest Biotech; catalog # EB11012) | 1:1000 |
|  | 488 Donkey anti-Goat (Jackson ImmunoResearch; catalog # 705-545-147) | 1:500 |

**Table 2: List of the cannula used in the surgery.**

| **Experiment** | **Details** |
| --- | --- |
| Fiber photometry (OmFC) | **Fiber details:** OmFC_ZF1.25_200/245-0.37_4.0mm_FLT_4.0mm  1.25 mm ferrule diameter, 200 µM fiber diameter, 0.37 numerical aperture, 4mm fiber  **Fluid injecto**r: FI_OmFC-ZF_100/170_4.4mm **Tubing:** Tubing_BTCOEX_25G_clear  (Doric lenses, Quebec, Canada) |
| Single Cannula for CART microinfusion | **Guide cannula:** C315G, 26G, 4mm  **Internal cannula:** C315I, 33G, 4.25mm **Dummy cannula:** C315DC, 33G, 4mm  (P1 technologies, VA, USA) |
| Bilateral cannula for CART microinfusion | **Guide cannula:** C235G-2.0/SPC, 26G, 2.25mm  **Internal cannula:** C315I/SPC, 33G, 2.6mm **Dummy cannula:** C235DC/SPC, 33G, 2.25mm  (P1 technologies, VA, USA) |
